# *Gouania willdenowi* is a teleost fish without immunoglobulin genes

**DOI:** 10.1101/793695

**Authors:** Serafin Mirete-Bachiller, David N. Olivieri, Francisco Gambón-Deza

## Abstract

In the study of immunoglobulin V genes in fish genomes, we found that the species *Gouania willdenowi* does not possess any such regions, neither for the heavy chain nor for the light chains. Also, genes that code for the immunoglobulin constant regions were also not found. A detailed analysis of the chromosomal region of these genes revealed a deletion in the entire locus for regions of the heavy and light chains. These studies provide evidence that this species does not possess genes coding for immunoglobulins. Additionally, we found the genes that code for CD79a and CD79b protein molecules have also been deleted. Regions for the T*α*/*β* lymphocyte receptors are present but the T *γ*/*δ* receptors were not found. In transcripts of two other Gobiesocidae species, *Acystus sp.* and *Tomicodon sp.*, no antibody sequences could be detected, possibly indicating the absence of immunoglobulins in all species of this family.

## 1. Introduction

The genes that code for immunoglobulins (IG), T lymphocyte receptors (TCR), and major histocompatibility class I and II complexes (MHC-I and MHC-II, respectively) had their origins in jawed vertebrates. In the case of IG, a locus was generated for the immunoglobulin heavy chain (IGH) genes and another independent locus for the light chains. Once originally constituted in these early species, these loci have remained intact throughout the evolution of all vertebrates.

The IG loci of teleosts has an organization similar to that found in terrestrial vertebrates. The locus for the IG constant regions (IGC) contains a wide region for the variable heavy-chain (VH) genes, the presence of JH and DH minigenes, and genes for IgM and IgD located downstream. An additional gene has been found in some species for the constant chain, referred to as IgT. These genes code for antibodies that may be in soluble form or as membrane receptors for B cells.

The B-lymphocyte receptor (BCR) is necessary for the generation of antibody response. Both in mammals and fish, it is a multimeric molecular complex formed by surface Ig and two other non-covalently associated proteins (CD79a and CD79b), which are necessary for receptor expression and function. Both CD79a and CD79b have an extracellular immunoglobulin-like domain, a transmembrane domain, and a cytoplasmic stem with an ITAM motif essential for signal transduction (Michael, 1995). In mammals, the CD79a molecule is encoded by the MB-1 gene (Sakaguchi et al., 1988), and the CD79b molecule by the B29 gene (Hermanson et al., 1988). Gene orthologs have been described in fish (Liu et al., 2017).

The Gobiesocidae family (clingfishes) belongs to a clade of teleosts (Ovalentaria). Ovalentaria includes 44 families with more than 5000 species. This term derives from the Latin where ‘ova’ means egg and ‘lenta’ means sticky. It is estimated from phylogeny studies that its appearance was approximately 91 million years ago (MYA), determined by analyzing 10 nuclear genes of 579 species (Near et al., 2013). Clingfishes have a small size (<8 cm), flattened body, reduced swim bladder, and an adhesive ventral disc that allows them to join surfaces (Briggs, 1955). They are distributed widely across the world in more than 170 species and 50 genera (Conway et al., 2018), having representatives both in the Pacific and Atlantic oceans, as well as in tropical and temperate seas. Recently, Gobiesocidae has been included as one of the 17 families that are part of the cryptobenthic reef fishes (CRF), in which at least 10 percent of the species of a family are less than 5 cm (Brandl et al., 2018).

*Gouania willdenowi* (blunt-snouted clingfish), has a small size (5 cm max.) and usually inhabits the interstices of gravel beaches. It is endemic to the Mediterranean Sea and its distribution ranges from the Eastern to Western Mediterranean. It belongs to the order of the Gobiesociformes (specifically, to the Gobiesocidae family), being the only representative of its genus, however recently five species have been described using morphometric studies and analysis of different genes (Wagner et al., 2019).

In this article we present surprising findings that suggest that the clingfish, *Gouania willdenowi*, lacks immunoglobulins as well as CD79a and CD79b. Therefore, this is a fish without BCRs and without essential and central elements of the adaptive immune response, such as antibodies.

## 2. Methods

Genome sequences were obtained from the GeneArk database repository (https://vgp.github.io). These datasets constitute complete high quality assemblies without gaps. Gene transcripts of *Gouania willdenowi* (RNA-seq) were obtained from the NCBI and correspond to the files ERR2639611, ERR2639610 and ERR2639609. The gene transcripts of fish deposited in China National Genbank were also studied from sequences reported from the Transcriptomes of 1,000 Fishes (FISHT1K) project.

The V regions of immunoglobulins and T lymphocyte receptors were identified from the genome datasets by the Vgeneextractor program (Olivieri et al., 2013). This program scans the entire genome and obtains most of the V regions. When the entire genome is scanned, a large number of false positives is obtained. To eliminate them, two filtering procedures were used. First, each sequence obtained by the Vgeneextractor was scored by predictions from a deep learning neural network (using Tensorflow) that was trained from known sequences of the 7 loci present in vertebrates (IGHV, IGKV, IGLV, TRAV, TRBV, TRGV and TRDV). A selection threshold on the predicted score was used to retain a sequence in the positive set. Subsequently, a second procedure was used to decide which V gene sequences that were consecutive in genomic segments were selected, whereas those at isolated sites in the genome were eliminated (i.e., probable false positives).

Specific searches for genes coding for the immunoglobulin constant regions and B-cell receptor genes (CD79) were performed with known sequences of *Oreochromis niloticus* (Ovalentaria, (Wu et al., 2019)). Additionally, from the annotations of the genomes of the NCBI, genes surrounding the target genes were studied carefully.

The divergence time of the species in this study was determined by the online evolutionary timescale tool at http://www.timetree.org/, which we also used to construct the species tree.

For sequence alignments, the SEAVIEW (Gouy et al., 2009) and Jalview (Waterhouse et al., 2009) programs were used together with the clustal*ω* algorithm (Sievers & Higgins, 2014). Phylogenetic trees of amino acid sequences were constructed from phyML (Guindon et al., 2010) using default parameters within SEAVIEW. Graphic representations of trees were made with Figtree (Rambaut A, 2019).

The exon search for MHC-I and MHC-II antigens was performed with our Python-based software MHCfinder (Olivieri & Gambon-Deza, 2018). This application uses machine learning predictions to identify sequences of the three main MHC-I exons (i.e., exon-2, 3 and 4), the two main MHC-II alpha-chain exons (i.e., exon-2 and 3), and the MHC-II beta-chain exons (i.e., exon-2 and 3). Graphic representations of the exon maps were constructed with the Python library, Genomediagram (Pritchard et al., 2005).

## 3. Results

With Vgeneextractor, V regions were searched for fish species with completely sequenced genomes without gaps (from the GeneArk public repository https://vgp.github.io/, described in Methods) (Table 1). While all teleost fishes studied have an IGHV locus and several loci similar to the IGKV regions of terrestrial vertebrates, we did not find neither the VH nor the VK regions in the two *Gouania willdenowi* genomes studied. Also, IG constant region (IGC) genes were not found in specific searches in these genomes. However, TRAC and TRBC genes were found, while TRGC and TRDC were not. Finally, the genes for CD79a and CD79b molecules that conform the B cell receptor were not found.

**Table 1:** Number of V regions per locus

| Specie | ighv | igkv | iglv | tr(a/d)v | trbv | trgv |
| --- | --- | --- | --- | --- | --- | --- |
| A._centrarchus | 240 | 93 | - | 87 | 97 | 15 |
| A._percula | 54 | 13 | - | 24 | 30 | 4 |
| A._testudineus | 203 | 29 | - | 84 | 72 | 22 |
| C._gobio | 36 | 5 | - | 12 | 40 | - |
| D._cupleoides | 87 | 29 | - | 49 | 26 | 7 |
| E._calabaricus | 76 | 105 | - | 55 | 41 | 9 |
| E._lucius | 67 | 80 | - | 54 | 41 | 11 |
| E._naucrates | 58 | 97 | - | 42 | 44 | 14 |
| G._willdenowi | - | - | - | 77 | 57 | - |
| L._chanluniae | 37 | 1 | - | 107 | 71 | 13 |
| M._armatus | 145 | 144 | - | 115 | 41 | 13 |
| P._flavescens | 107 | 47 | - | 29 | 53 | 1 |
| P._ranga | 86 | 37 | 4 | 54 | 93 | 23 |
| S._aurata | 302 | 102 | - | 274 | 299 | 64 |
| S._formosus | 164 | 193 | - | 154 | 62 | 19 |
| S._trutta | 75 | 49 | - | 131 | 218 | 9 |
| T._rubripes | 78 | 21 | - | 55 | 27 | 18 |

We analyzed the microsynteny of this regions in Ovalentaria from NCBI genome datasets with an assembly at the chromosomal level. In all the fishes studied, CD79a is flanked by the LIPE and AERHGF1 genes. In *Gouania willdenowi*, no CD79a gene is found between these two genes (Figure S1). Similarly, CD79b is always found between SCN4A and IFNA3-like genes in other fishes, but is absent in the case of *Gouania willdenowi* and there is a inversion (Figure S2). Genes for IGH (heavy chains) and IG light chains in *Gouania willdenowi* are not located among those flanked genes in other Ovalentaria species (Figures S3, S4)

To identify the moment in evolution when the loss of these genes occurred, we studied transcriptome sequences from the Fish-T1k project (Sun et al., 2016). Since *Gouania willdenowi* is in the genus Ovalentaria, we performed of BLAST search of IgM of all Ovalentaria transcripts. All fishes from this genus have IgM sequences except for two species, *Tomicodon sp.* and *Acyrtus sp.*. As *Gouania willdenowi*, these two species belong to the Gobiesocidae family, suggesting that immunoglobulins are absent from species of this family. The estimated divergence time of 77 MYA between the Gobiesocidae family from Blenniiformes, the next closest in evolution, may be the origin of this genetic modification. In the three Gobiesocidae species studied, the sequences for CD79a and CD79b and for TRGC and TRDC were not found. However, the V*α* and V*β* chains genes (TRAV and TRBV, respectively) of TCR were found.

The sequences of the V genes for the *α* and *β* chains, TRAV y TRBV respectively, were studied in detail in *Gouania willdenowi*. From Vgenextractor (see Methods), 77 TRAV exon sequences were obtained from scaffold 258 and in the Super Scaffold 5; while 57 TRBV exon sequences were found in the Super Scaffold 7. The TRBV exon sequences display similar variability to that found in other species with 8-10 clusters from similar regions (Figure 2 and Supplementary Figure S6). The TRAV exon sequences are quite similar to each other with particularities in the sequence (Figure 2 and Supplementary Figure S5). In these TRAV exons, the framework regions are highly conserved; in all the exons there is a cysteine in the L2 region at the beginning of the sequence, and another cysteine in most of the sequences in the third framework region (in addition to the canonical cysteine).

**Figure 1:**
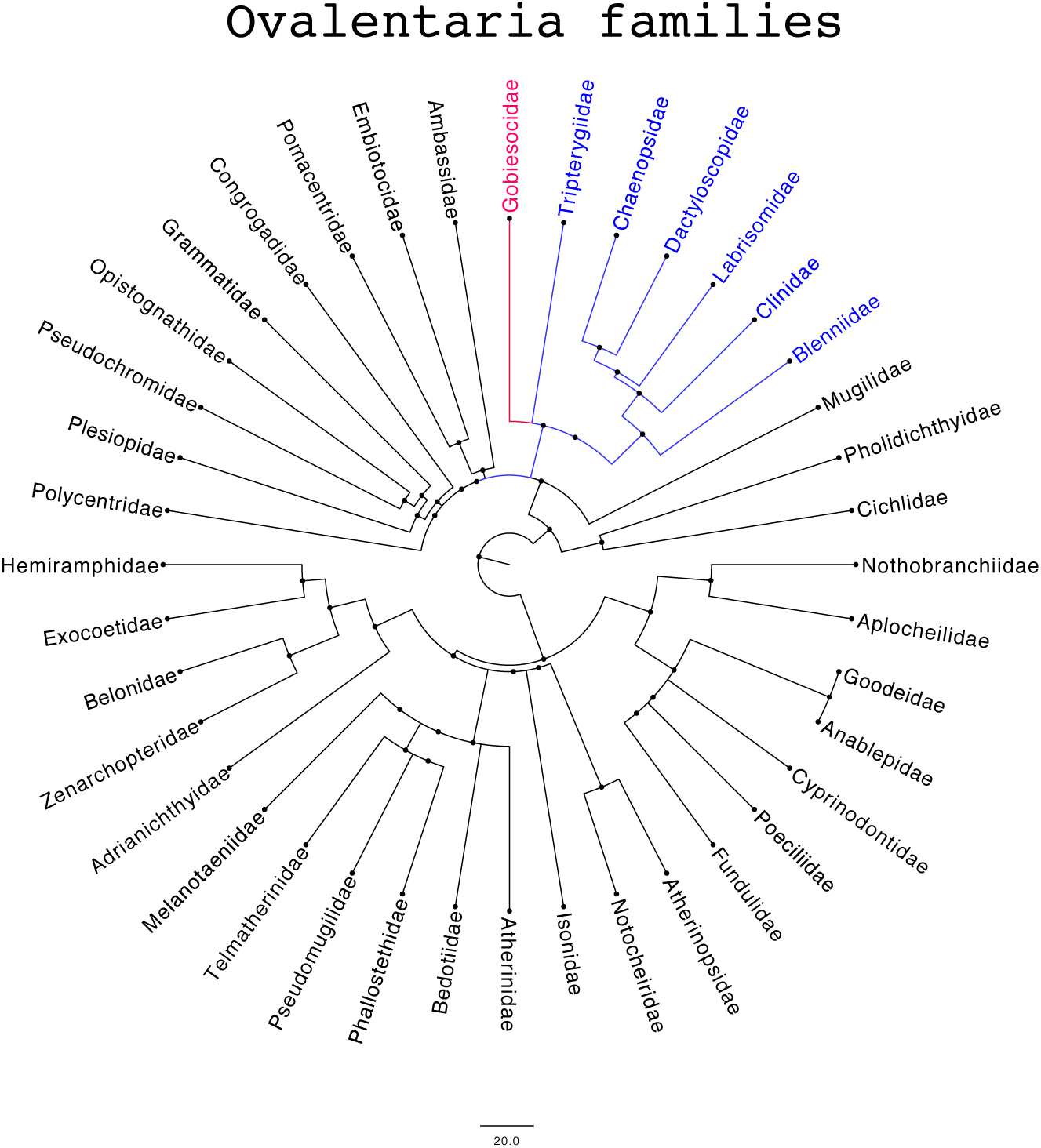
The phylogenetic tree of the Ovalentaria families of teleost fish. The tree was constructed from evolutionary data provided by Timetree (http://www.timetree.org/). The clade is indicated corresponding to the Blenniiformes within which is the Gobiesocidae family, of which *Gouania willdenowii, Acyrtus sp. and Tomicodon sp.* belong.

**Figure 2:**
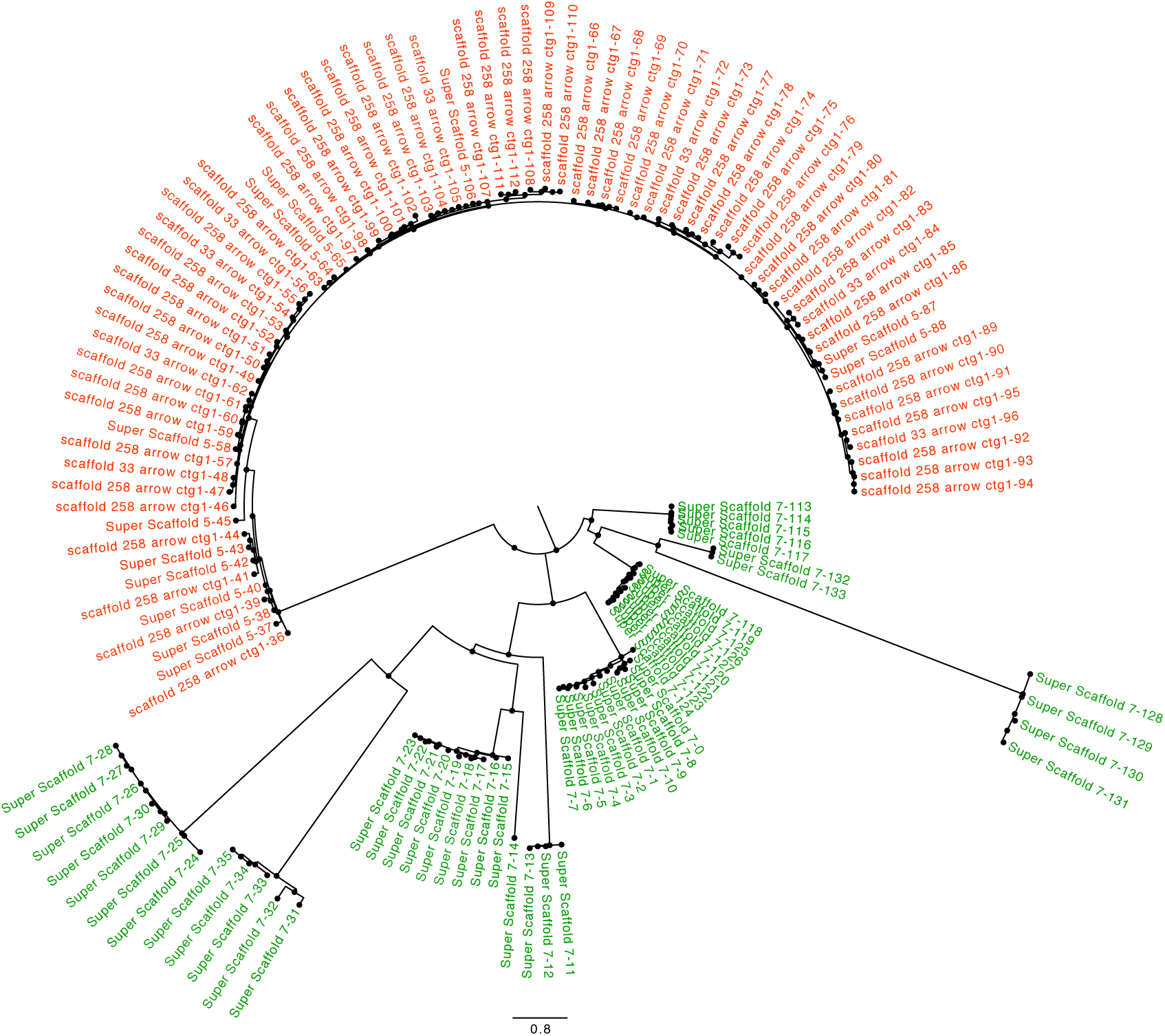
The phylogenetic tree of the V exons amino acid sequences from *Gouania willdenowi*. The exon V sequences were aligned with the ClustalΩ, tree construction with PhyML with the LG matrix. The sequences of exons V*β* are displayed in green, while those for V*α* are in red.

This study also searched for the presence of MHC-I and MHC-II genes. These genes are related to the adaptive immune response and the loss of MHC class II genes has been described in cod species. With MHCfinder (see Methods section), which uses machine learning to recognize the main MHC exons, we found 31 genes that are probably viable for MHC-I and one gene for an MHC-II alpha chain and another for an MHC-II beta chain (Figure 3). Unlike that found in mammals, these MHC-I genes are found on chromosomes different from those of MHC-II, which was already described in teleost fish (Kaufman, 2018).

**Figure 3:**
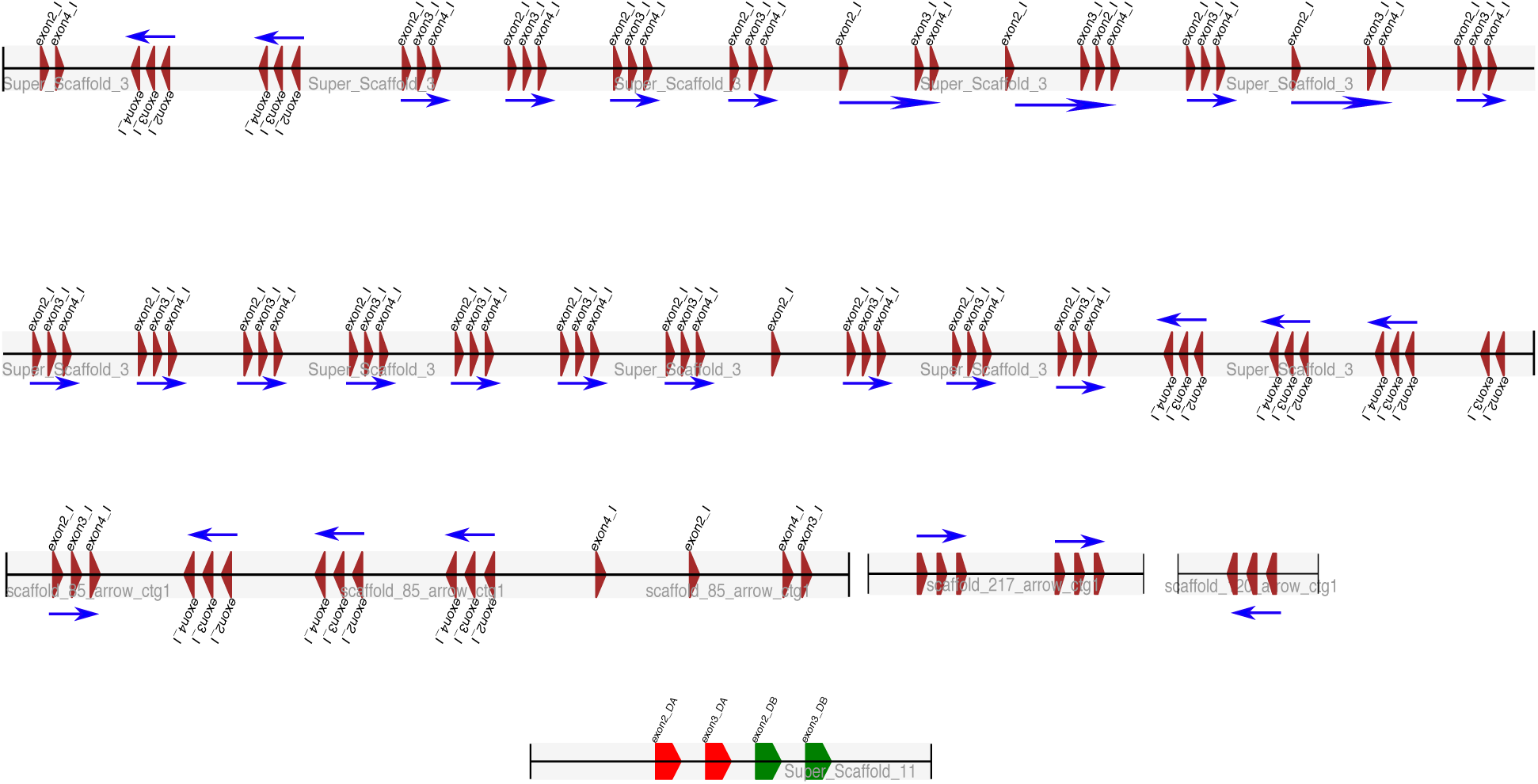
Schematic representation of the MHC-I and MHC-II exons found with MHCfinder in the genome of *Gouania willdenowi*. Exons that make up an MHC gene are grouped together based upon a simple heuristic rule, based upon their distance along the genome: they are grouped if the distance between them is less than 3000 kb, otherwise they are considered part of other genes or non-viable isolated exons. Super Scaffold 3 has most of the MHC-I genes, although they are also found in other segments of the genome. The MHC-II genes are located in the Super Scaffold 11.

## 4. Discussion

Due to the recent availability of complete fish genomes, we could undertake a complete study of IG and MHC genes in these species, allowing us to discover the absence of immunoglobulin genes in *Gouania willdenowi*. This is the first such case described in vertebrates.

Apart from the absence of IG genes, the CD79a and CD79b genes that make up the B lymphocyte receptor are also absent, suggesting they were lost first in evolution. It is difficult to imagine that the loss of antibody genes occurred in the initial evolutionary process since several loci are located on different chromosomes. The loss of CD79 may explain the subsequent disappearance of the IG genes. The absence of the CD79 proteins that are necessary to form the B-lymphocyte receptor, left the immunoglobulins without environmental selection, subsequently leading to the deletion of these IG genes over time.

We found that the loss of these IG genes is restricted to Gobiesocidae. The three representative fish of this family that we studied all lack these genes and neither CD79 nor the IGs are present in the published transcriptomes. The loss of the T *γ*/*δ* receptor is more difficult to explain. This receptor is not present in Squamata reptiles (Olivieri et al., 2014), suggesting that it is expendable in evolutionary lines of jawed vertebrates. Based upon the data available, we cannot discern whether the loss of this receptor in the Gobiesocidae family is linked to the loss of IG genes, or corresponds to another additional event.

*G. morhua* (Atlantic cod) has lost the genes for MHC-II, CD4, and CD74, thereby deprived of the classical adaptive immunity pathway to respond against bacterial and parasitic infections. Nonetheless, the Atlantic cod does not seem more vulnerable to infections. A probable explanation for this is that the development of compensatory mechanisms arising from an expansion in the number of genes that code for MHC-I (something that also observed in *Gouania willdenowi*) and a specific repertoire of TLRs for the Atlantic cod (Star et al., 2011). Similarly, a fish that is phylogenetically distant, such as *S. typhle* (pipefish), lacks MHC-II, the CD4 receptor, and TCRGD (Haase et al., 2013). All these facts suggest deletions of basic genes for immune response can occur in fish and these species can still be viable. We have not found alternatives to the absence of IG in *Gouania willdenowi*. The presence of at least 31 MHC-I genes is not considered exceptional in fish. Likewise, the peculiarities of the sequences of the TRAV (V*α*) are of uncertain significance, and the search for Toll-like genes does not give different results from those in other fish.

The absence of these genes at the transcriptome level were studied in *Tomicodon sp.* and *Acyrtus sp.* and were found to have similar results with respect to these genes as that of *Gouania willdenowi*. It is possible, that by extension of this evidence, that these genes are absent across the entire Gobiesocidae family. This would be surprising and would require further studies since this family is composed of more than 170 species worldwide with 50 genders.

Despite being unable to produce antibodies, it was demonstrated that *Rag1*^−/−^ *Danio rerio* (zebrafish) reach adulthood without obvious signs of infection under non-sterile conditions (Wienholds et al., 2002). Recently, it was shown that these *Rag1*^−/−^ fish show signs of early aging and a decreased lifespan when compared to wild-type fish (Novoa et al., 2019). In humans, agammaglobulinemia pathology is described by a mutation in the CD79 gene, which is a serious disease that forces patients to be treated with exogenous immunoglobulins. The fact that these fish can survive in nature suggests that the functionality of their immune system is different from that of mammals, at least in response to infection.

The main question raised by this study concerns the viability of a vertebrate without genes for antibodies. Antibodies were described in mammals as a defense mechanism against infection. Subsequently, autoimmune phenomena were discovered that indicate that their own structures can be recognized. In recent years, multiple immune responses to cancers or senescent cells are being described. Therefore, such studies indicate that the activities of the immune system are greater than those that were initially deduced. However, the extrapolation of what is found in mammals to fish does not have to be absolute. The acquired immune system was created with the first jawed vertebrates. In general, its creation has been attributed as a defense mechanism against infection, and therefore it can be complementary or redundant to the innate system already present in previous evolutionary lines.

Considering the existence of fish that do not possess antibodies yet are nonetheless viable species, suggests that the adaptive immune system may have been created primarily for functions other than infection, perhaps as a cellular control system to prevent the development of cancers or senescence. This immune system must have evolved in animals that moved to land as demonstrated by the appearance of new antibody genes with more effective class switch recombination (CSR) phenomena and affinity maturation processes. These adaptations probably extended the functionality of the immune system towards infection, which may explain the differences in viability between fish and subsequent species. This hypothesis could explain the flexibility observed, where components of the adaptive immune system are absent in some species of fish without their viability being compromised, while in mammals a severe immunodeficiency is generated. It is expected that as more fish genomes are available these questions can be explored further.

## Supplementary Material

**Figure S1:**
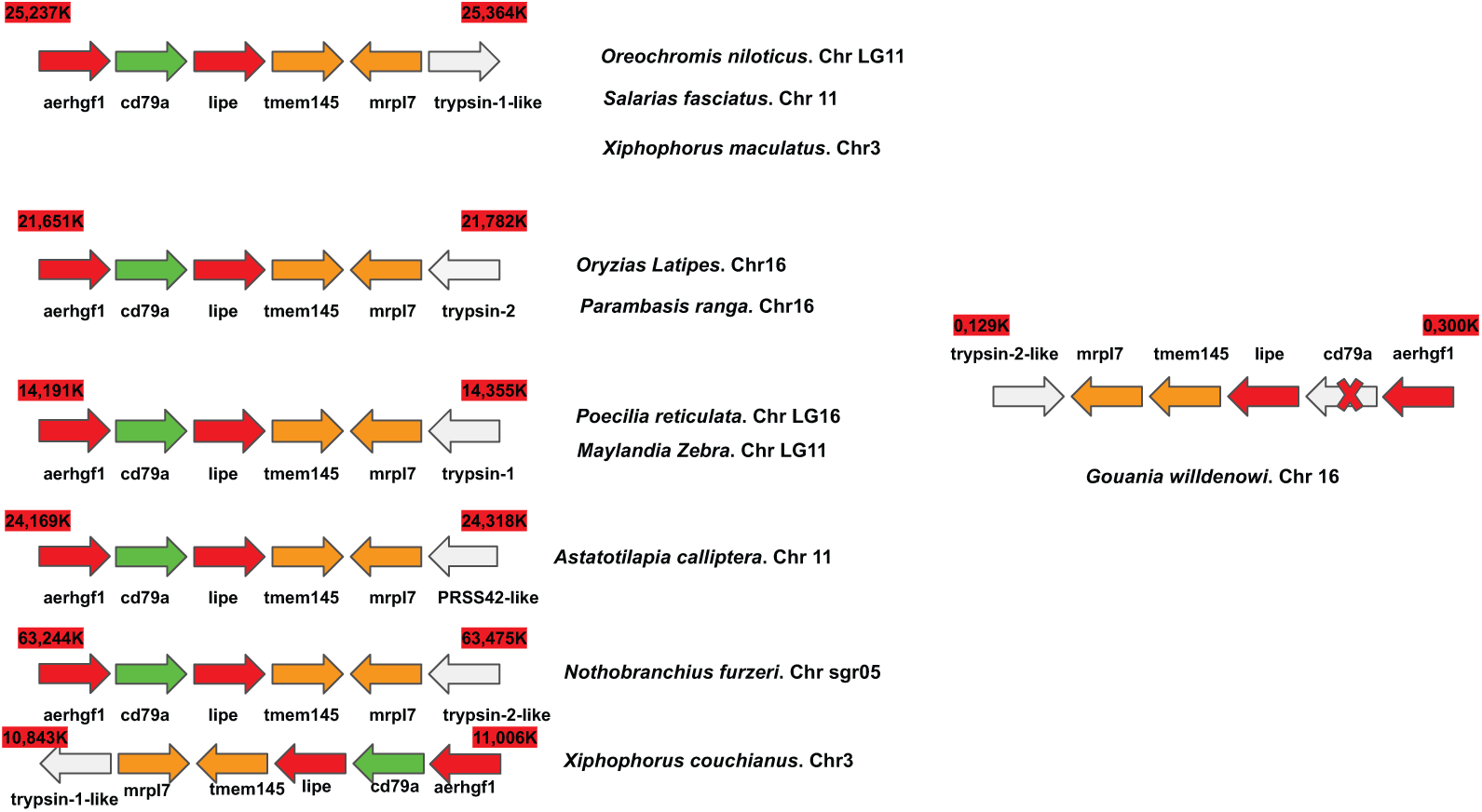
Gene microsynteny of CD79a identified in Ovalentaria fishes. The color coding is as follows: CD79a (green), genes flanking CD79a (red), other genes next to flanking genes (orange), different genes in Ovalentaria fishes (grey). In all Ovalentaria studied, CD79a is flanked by the AERHGF1 and LIPE genes except in *Gouania willdenowi*, where it is absent.

**Figure S2:**
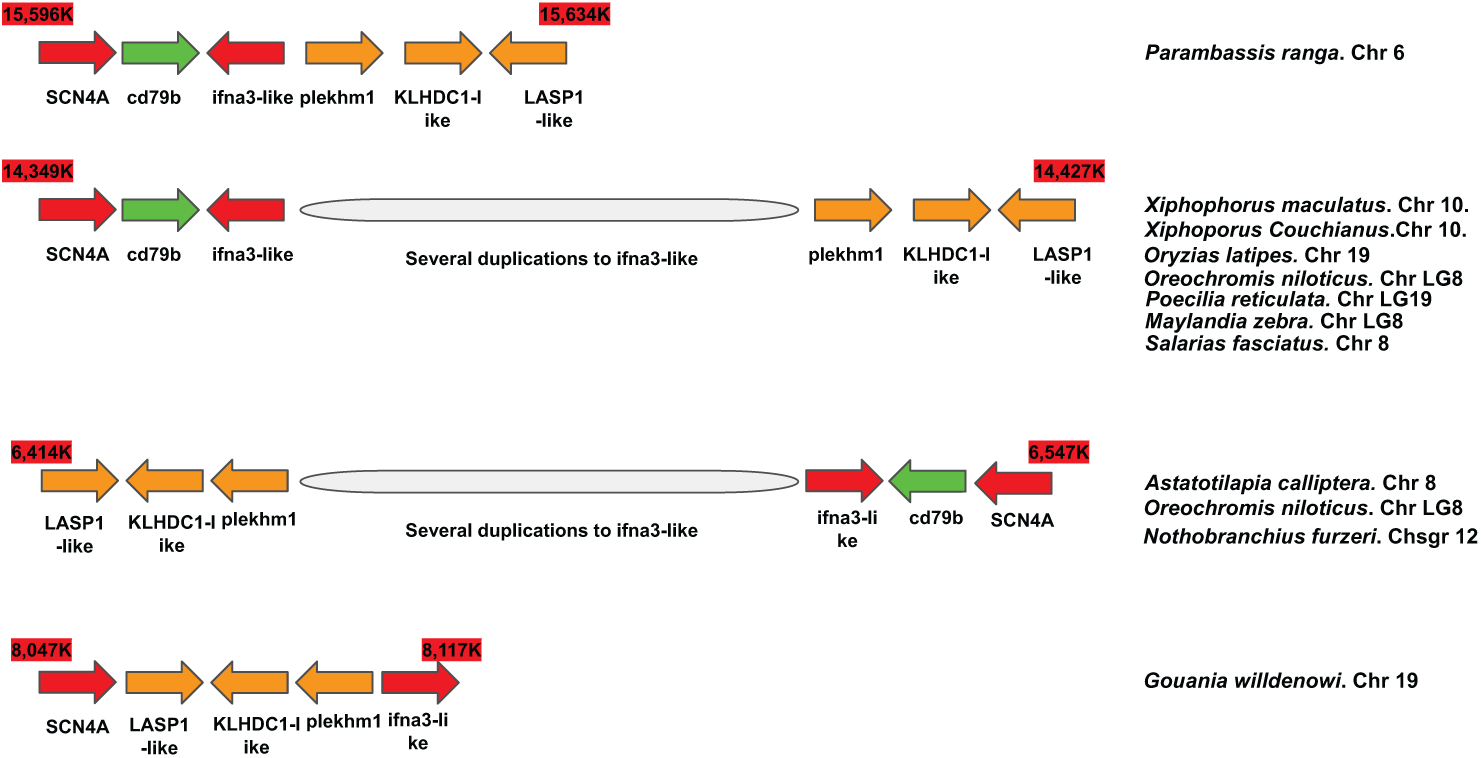
Gene microsynteny of CD79b identified in Ovalentaria fishes. The color coding is as follows: CD79b (green), genes flanking CD79b (red), other genes next to flanking genes (orange), different genes in Ovalentaria fishes (grey). In all Ovalentaria studied, CD79b is flanked by the SCN4A and IFNA3-like genes except in *Gouania willdenowi*, where it is absent.

**Figure S3:**
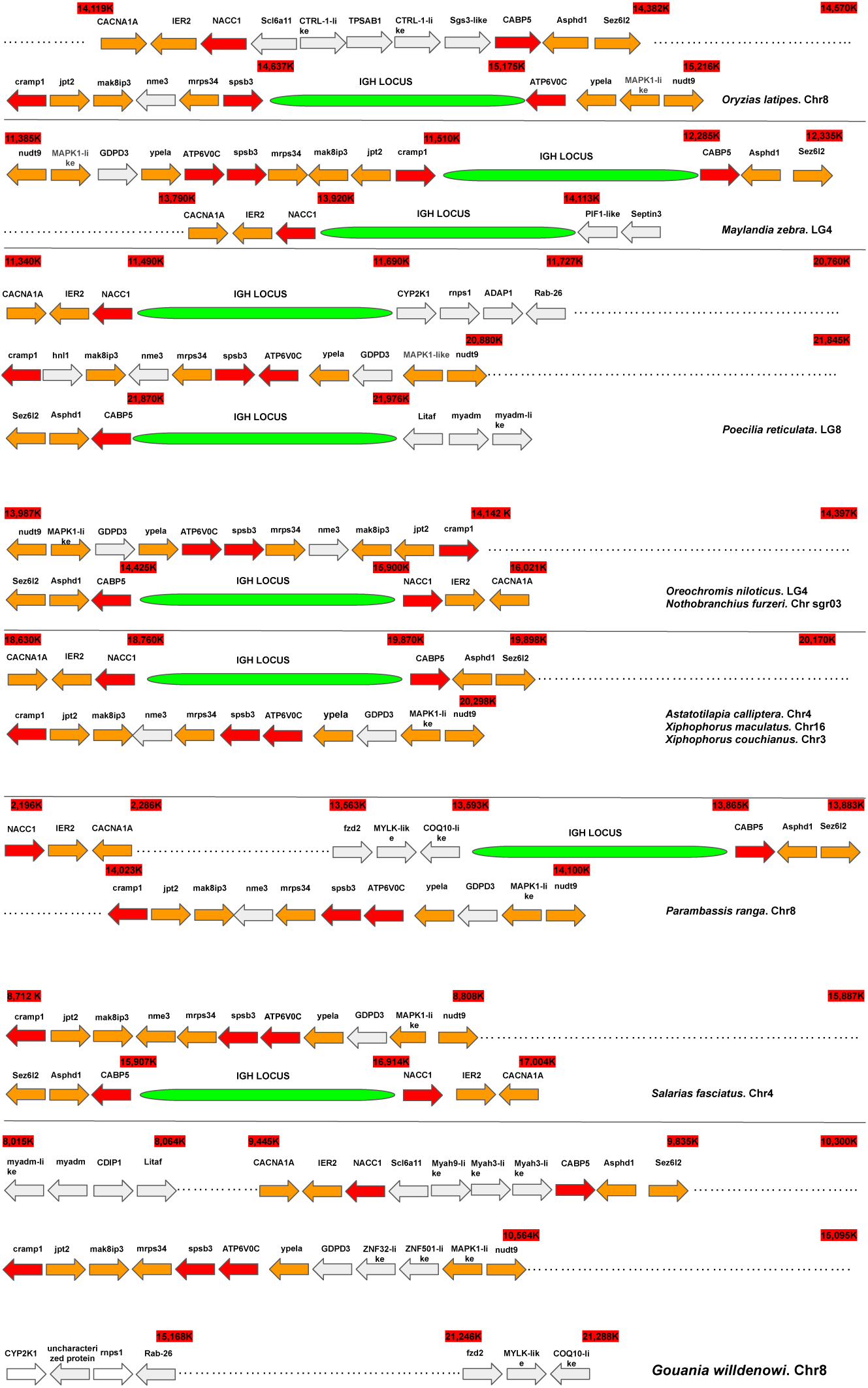
Gene microsynteny of the IGH loci identified in Ovalentaria fishes. The color coding is as follows: the IGH locus (green), genes flanking the IGH locus (red), other genes next to flanking genes (orange), different genes in Ovalentaria fishes (grey). The IGH locus is flanked by at least one of the following genes, SPSB3, ATPV0C, NACC1, CRAMP1 or CABP5 genes, all of which are present in Chr 8 of *Gouania willdenowi* but not in the IGH locus.

**Figure S4:**
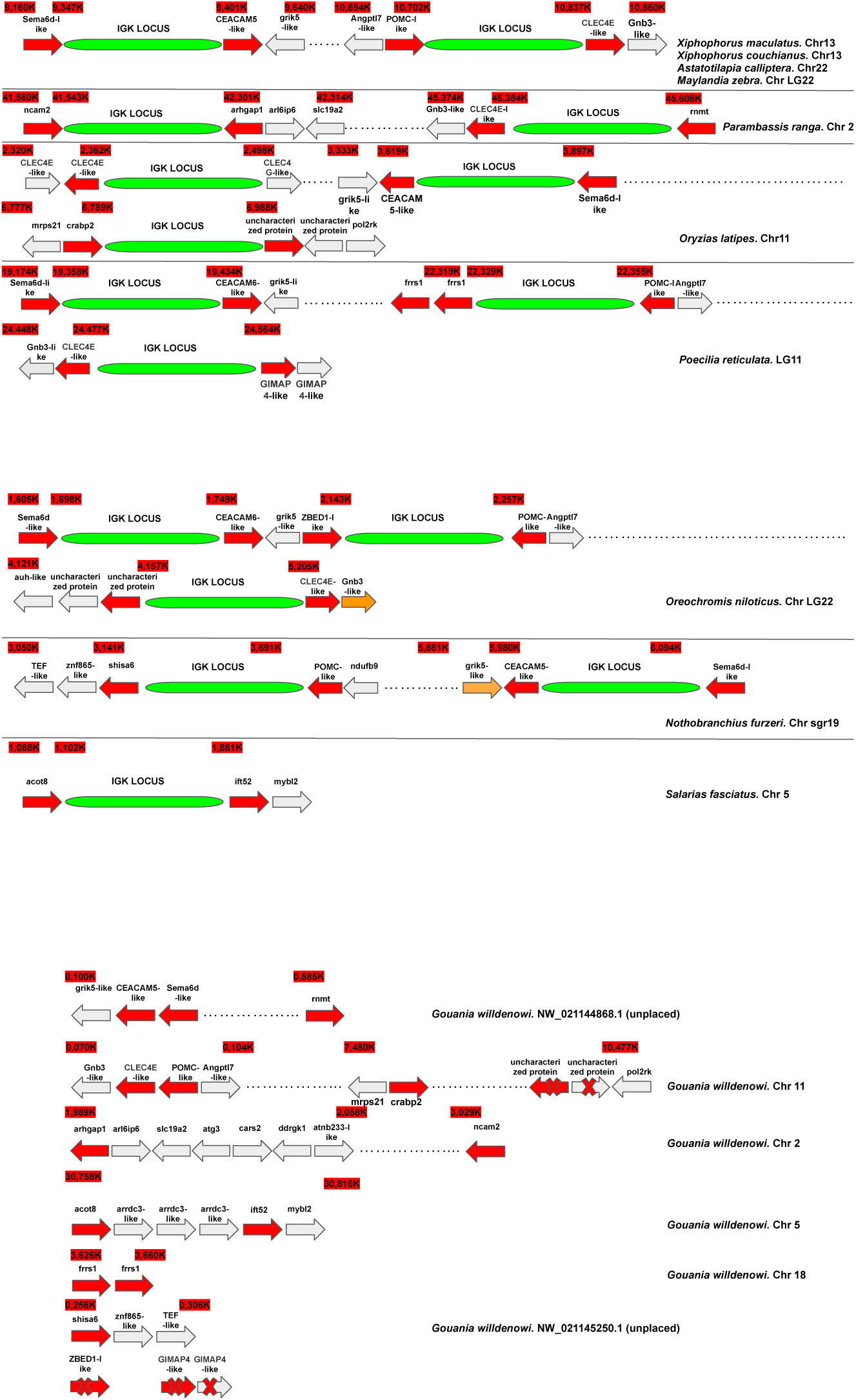
Gene microsynteny of the IGK loci identified in Ovalentaria fishes. The color coding is as follows: the IGK locus (green), genes flanking the IGK locus (red), different genes in Ovalentaria fishes (grey). The IGK is not observed in any case of *Gouania willdenowi*.

**Figure S5:**
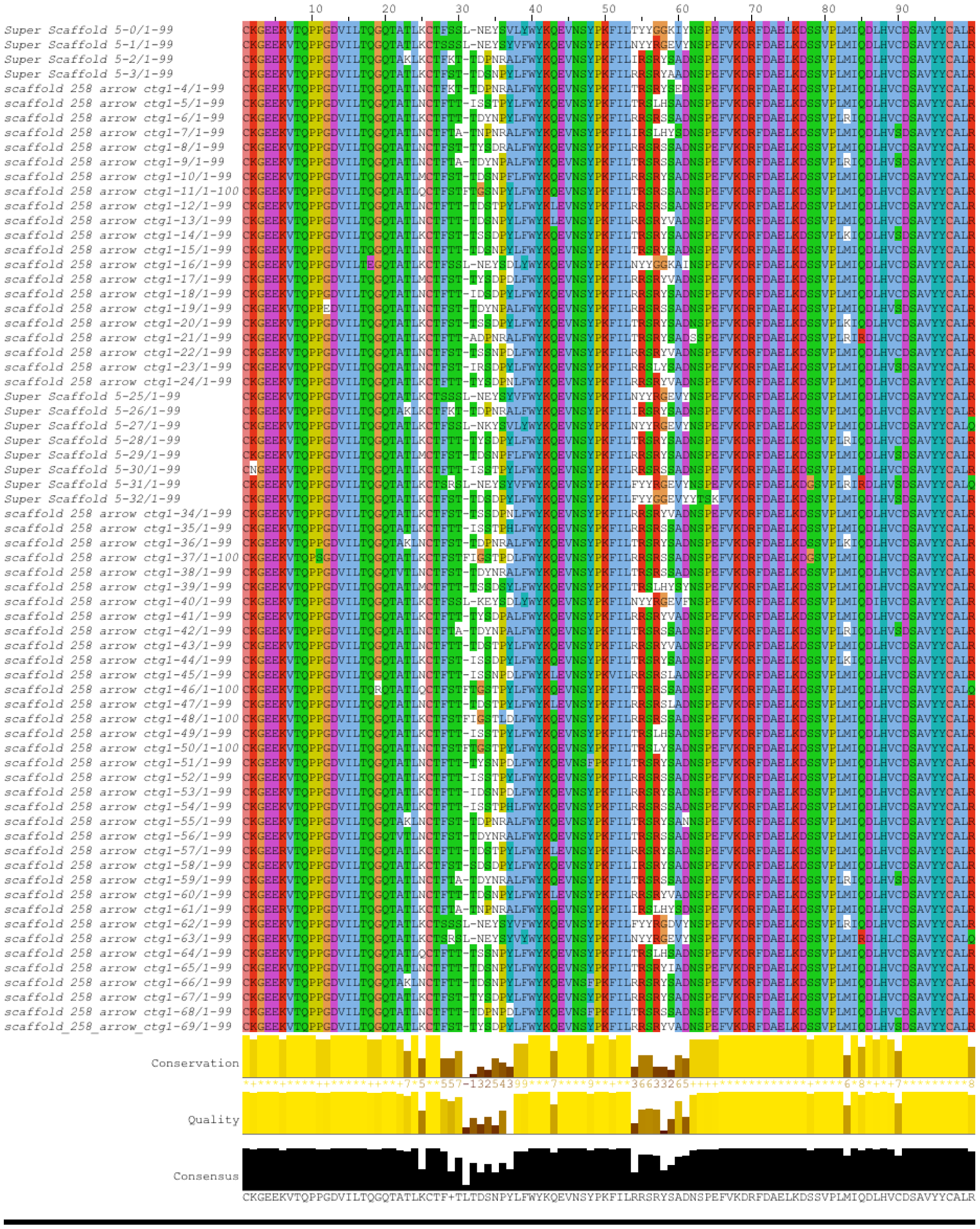
Deduced amino acid sequences of the sequences of V*α* exons (TRAV) obtained with Vgenextractor (see Methods). The graph was obtained with the Jalview. Notable are the conserved Framework regions, the presence of cysteine as the first amino acid, and the presence of an additional cysteine in the Framework 3 region.

**Figure S6:**
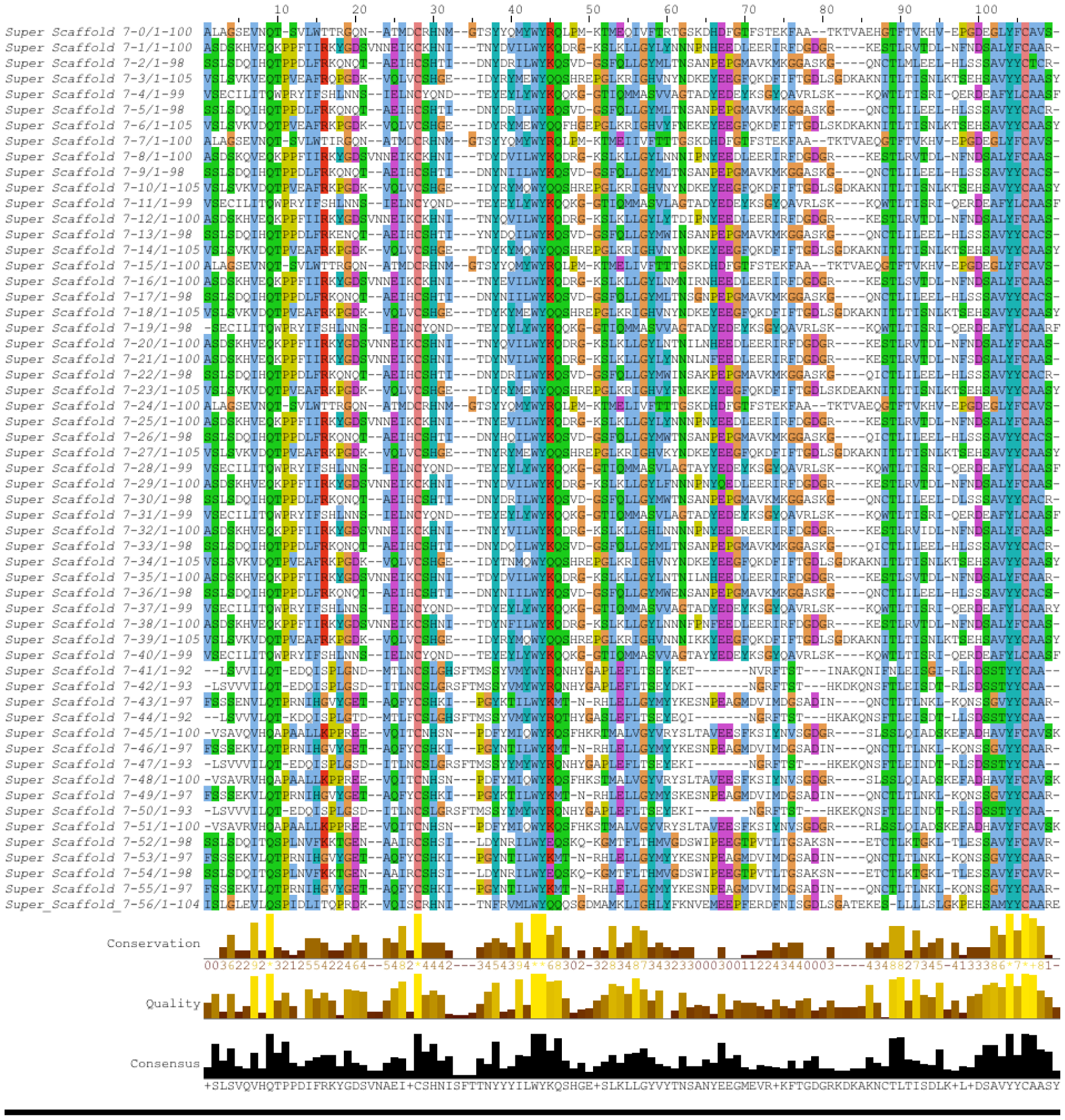
Deduced amino acid sequences of the sequences of V*β* exons (TRBV) obtained with Vgenextractor (see Methods). The graph was obtained with the Jalview.

